# Predicting the Three-dimensional Structure of the c-*KIT* Oncogene Promoter and the Dynamics of Strongly Coupled Guanine-Quadruplexes

**DOI:** 10.1101/2023.02.23.529733

**Authors:** Emmanuelle Bignon, Angelo Spinello, Tom Miclot, Luisa D’ Anna, Cosimo Ducani, Stéphanie Grandemange, Giampaolo Barone, Antonio Monari, Alessio Terenzi

## Abstract

Guanine-quadruplexes (G4s) are non-canonical DNA structures that play important protective and regulatory roles within cells, influencing, for instance, gene expression. Although the secondary structure of many human G4s is well characterized, in several gene-promoter regions multiple G4s are located in close proximity and may form three-dimensional structures which could ultimately influence their biological roles. In this contribution, we analyze the interplay between the three neighboring G4s present in the c-*KIT* proto-oncogene promoter, namely WK1, WSP and WK2. In particular, we highlight how these three G4s are structurally linked and how their crosstalk favors the formation of a parallel structure for WSP, differently from what observed for this isolated G4 in solution. Relying on all-atom molecular dynamic simulations exceeding the μs time-scale and using enhanced sampling methods, we provide the first computationally-resolved structure of a well-organized G4 cluster in the promoter of a crucial gene involved in cancer development. Our results indicate that neighboring G4s influence their mutual three-dimensional arrangement and provide a powerful tool to predict and interpret complex DNA structures that ultimately can be used as starting point for drug discovery purposes.

## INTRODUCTION

The presence of non-canonical DNA structures in the human genome, which may differ considerably from the well-known double helical arrangement, is nowadays established.^1,2^ Among such motifs, guanine quadruplexes (G4s) hold a special significance, and have been the subject of a high number of studies encompassing molecular, structural, and cellular biology.^3,4^ Not only the presence of G4s has been proven in living cells, but their crucial biological role has been stressed out. G4s are formed in guanine-rich regions of the DNA (and RNA) and are located in crucial, often non-translated, segments of the genome. In particular, they accumulate in the terminal regions of the chromosomes, i.e. the telomeres, where they exert a protective role to assure genome integrity.^5^ Telomeric G4s inhibit the reverse transcriptase activity of the telomerase, hence contributing to avoiding the development of an immortal phenotype, which is commonly found in cancer cells. As such, telomeric G4s are emerging as target of choice for the development of original chemotherapy strategies.^6^ Furthermore, G4s have been identified in gene promoter regions, and their stabilization has been correlated to the regulation of the corresponding genes.^7^ Depending on their localization upstream or downstream of the coding gene, G4 may activate or repress DNA transcription.^3,4^ Interestingly, G4-compatible sequences have been identified in the promoter regions of a number of genes related to cancer development and progression (e.g., c-*MYC*, c-*KIT* and k*RAS*)^7^ In general, oncogenes harbor more G4s than tumor suppressor genes, further justifying the exploitation of their stabilization (to repress their transcription) via small ligands as a possible chemotherapeutic strategy.^4^ It is worth pointing out that G4s have also been identified in the genome of bacteria,^8^ including antibiotic-resistant pathogens, and in different viruses.^9,10^ As for the latter point, G4s play an important regulatory role in the lifecycle of different viral agents,^9,10^ including retroviruses, such as human immunodeficiency virus (HIV),^11^ or emerging RNA viruses, such as SARS-CoV-2,^12–15^ Zika,^16^ and Tick-borne encephalitis virus.^17^

From a structural point of view, G4s are constituted by an ensemble of guanines organized in tetrads and coupled via Hoogsteen hydrogen bonds. The tetrads are further superposed via their stabilizing π-stacking and the favorable interaction with monovalent cations (especially K^+^) occupying the central channel. Such an arrangement gives rise to a very rigid core, which is however flanked by rather flexible loops of different length, connecting the Hoogsteen-bonded guanines.^18,19^ Despite the rigidity of their core, G4s exhibit different conformational topologies, depending on the relative orientation of the backbone between adjacent guanines: namely parallel, antiparallel, and hybrid arrangements. Interestingly, while the parallel arrangement is the preferred one in RNA G4s, within DNA G4 motifs are more polymorphic and populate all the three main topologies, depending on the specific sequence and the environmental conditions. Transitions between parallel and hybrid or antiparallel topologies may be induced by changes in the central metal ion, or the crowding of the medium.^20–23^ Noteworthy, G4s have been shown to be extremely resilient to the inclusion of DNA lesions, including oxidative lesions and abasic sites, and the recruitment of DNA-repair enzymes by damaged G4s has also been associated to epigenetic control.^24–26^

Despite the extreme importance of the characterization of the proper folding of single G4s from truncated sequences, for instance by X-ray, NMR and electronic circular dichroism (ECD) spectra,^27^ this is usually not sufficient to properly infer their behavior in the more complex system represented by DNA. As mentioned, in gene promoter regions G4s are usually coupled in clusters and this complex organization may lead to a well-defined three-dimensional of the individual G4s, which may strongly alter the interaction with their biological partners like helicases or with targeting molecules.^28–30^In a sense, and in analogy with protein or RNA classifications, while individual G4s may be assimilated to secondary structure elements, their interaction, especially when present on the same strand, could be defined as a tertiary arrangement.

A perfect example of how complex the mutual arrangement of multiple neighboring G4s could be is represented by c-*KIT* gene promoter which features three G4s in close proximity,^31^ namely WK1 and WK2 quadruplexes intertwined by the two tetrad WSP (see Figure 1), a system that was recently studied experimentally.^32^

**Figure 1.**
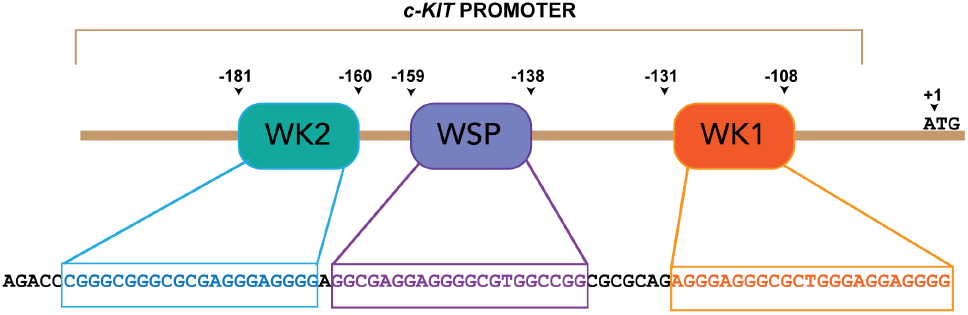
Schematic representation of the wild-type G4 units (WK2, WSP, and WK1) disposition in the *c-KIT* promoter.

Interesting, the isolated WK1 and WK2 sequences in solution fold into parallel quadruplexes,^33,34^ while the spectral signature of WSP is consistent with that of an antiparallel folding.^35,36^ However, the ECD of the complete sequence presents signatures compatible only with a parallel G4, suggesting that cooperativity is indeed favoring the latter arrangement for WSP. WK2 and WSP are suggested to form a superstructure of two coupled G4s.^32,37^ It is still unclear whether the cross-talk between these G4s actually promotes a conformational transition on WSP or simply inhibits the antiparallel folding. Overall, a clear atomistic picture of the ternary aggregate in the *KIT* promoter is missing. Moreover, the factors inducing the stabilization of the parallel arrangement of WSP instead of the antiparallel one remain elusive. A sub-system system comprising the linked WK2 and WSP has also been recently studied computationally, however with a focus oriented more towards the elucidation of the role of ions and bridging loops.^38^

In the present contribution, we use long-scale all-atom molecular dynamic (MD) simulations to fill this gap and provide the first computationally-resolved atomistic structure of a tertiary arrangement in a biologically relevant cluster of G4s. Furthermore, we will also identify the molecular bases driving the interplay between the individual G4s of the c-*KIT* promoter.

## RESULTS

A 78-bases-long single-stranded sequence including all three wildtype G4s in the c-KIT promoter has been reconstructed from the experimentally resolved structures of the individual WK1 and WK2 G4s, as detailed in the Methodology Section. Concerning the WSP motif, both topologies have been considered, i.e. parallel and antiparallel arrangements, the latter being the structure observed in solution by ECD. The results of the equilibrium MD simulation can be found in Figure 2, where the rather different behavior of the two arrangements is highlighted. In the case of WSP treated with its parallel arrangement in the context of the long sequence used, the three most populated cluster extracts from the MD trajectory suggest that a supramolecular tertiary organization is established in which WSP interacts strongly with the WK2 G4. Indeed, the close proximity of the two units and the similar parallel arrangement allows the formation of stabilizing interactions between the tetrads of the individual G4s, although an ideal π-stacking arrangement could not be obtained. This, in turn, leads to a relative short distance between the center of mass of WSP and WK2. The necessity of keeping the stacked conformation with WK2 is coupled with an increase of the distance with the third G4, WK1, whose center of mass is kept at more than 50 Å from WSP. Interestingly, a cross-talk between WK2 and WK1 with the formation of a stacked superstructure of the two G4s has been proposed by Rigo and Sissi,^37^ but this study does not mention whether WSP maintain its antiparallel nature at room temperature when part of the G4-G4 structure. In a previous work, some of us ruled out experimentally the antiparallel nature of WSP in the context of the long sequence.^32^

**Figure 2.**
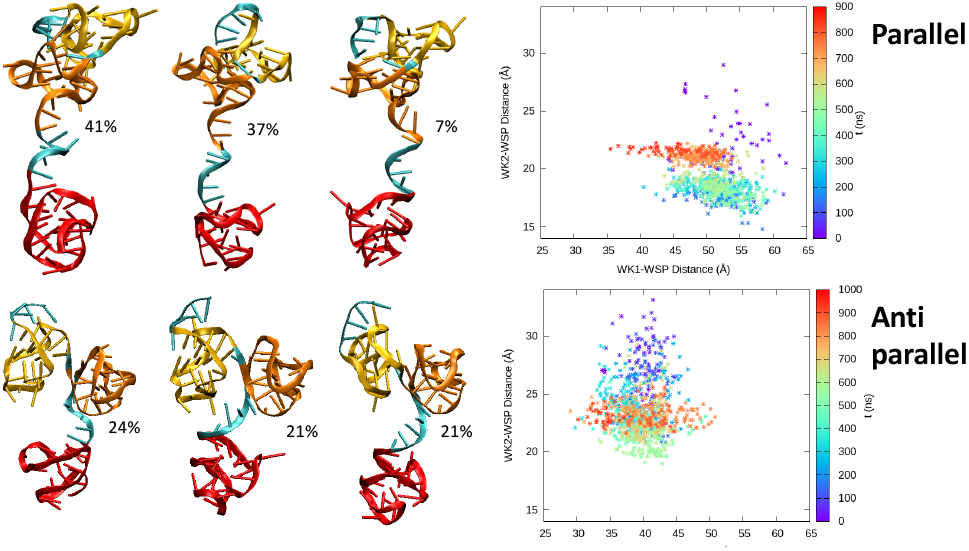
Structure of the three most representative conformations issued from the MD simulations of the c-*KIT* promoter sequence, featuring a parallel (top) or antiparallel (bottom) arrangement for WSP. WK1 is represented in red, WSP in orange, and WK2 in yellow, the weight of the corresponding structures in the MD simulation ensemble is also given. On the right, 2D plots of the time-evolution of the inter-G4 distances (centers of mass of WK1 and WSP1, and WK2 and WSP) correlation for the two topologies.

The stable stacked conformation between the two most proximal G4s is however broken when WSP assumes the antiparallel conformation, which is the favorable one observed for the isolated quadruplex in solution. As a consequence, and despite the fact that WSP and WK2 are separated by only one bridging nucleotide, the two G4s behave rather independently, as can be seen from the representative structures of the most populated clusters characterized by the simulation. In this respect, we notice that the higher flexibility of the WSP antiparallel-containing system is also reflected in a larger distribution of the accessible conformations, the three main structures harboring almost degenerate weights close to 21%. Interestingly, the three dominant conformations differ mainly by the relative orientation between WSP and WK2, as defined by the backbone dihedral angle of the bridging nucleotide. This orientational variability is also coherent with the enhanced flexibility and hence uncoupled behavior of the two G4s. When analyzing the time evolution of the distance between the couple WK2/WSP and WK1/WSP (see Figure 2), we may observe an opposite trend compared to the parallel situation. For the antiparallel WSP, the uncoupling with WK2 leads to a greater distance between the two G4s, which are no more interacting. On the contrary, the enhanced flexibility is translated into a stronger bending of the arrangement of the full sequence, hence into a shortening of the WK1/WSP distance which is now comprised in the range 30 - 40 Å,after 1 μs of simulation.

The results obtained from the equilibrium MD simulations clearly confirm the observation from ECD and melting assays recently reported in our previous work.^32^ Indeed, we may observe a cooperative behavior, and the establishment of favorable π-stacking interactions with WK2, only in the case of a parallel arrangement of WSP. On the other hand, the three G4s act independently when adopting the antiparallel conformation of the central member. These results show that the emergence of the tertiary arrangement between WSP and WK2 is acting as a stabilizing force. In turn, the intermolecular-driven stabilization of the aggregate contributes to the displacement of the equilibrium between antiparallel and parallel folding of WSP. This observation explains why, differently from the case of the isolated constituents, the ECD spectrum of the longer c-*KIT* sequence presents only spectral signatures associated with parallel arrangements.

To confirm the different behavior between the two WSP conformations, avoiding biases due by trapping in local minima of the free energy potential, we carried out a Gaussian Accelerated MD (GAMD) enhanced sampling. The corresponding corrected free energy profiles for the parallel and antiparallel settings are shown in Figure 3. Globally, all considerations issued from the equilibrium MD simulations are confirmed. In particular, when we consider the distance between the center of mass of the WK1/WSP and WK2/WSP couples, we clearly see that both arrangements present a single free energy minimum, indicative of a rather stable arrangement. However, the two systems also present interesting differences. The parallel WSP has a free energy minimum corresponding to a WK2/WSP distance of about 20 Å, and to WK1/WSP distance of about 60 Å. The corresponding representative snapshots evidence a stacking interaction between WK2 and WSP, observed also in the unbiased trajectories. On the other hand, in the case of the antiparallel conformation the WK2/WSP minimum energy distance is around 28 Å, and the WK1/WSP distance is reduced to 45 Å. The visual inspection of the snapshots corresponding to the minimum free energy clearly highlights the absence of the π-stacking interaction between WK2 and WSP and the rotation of the two G4s to assume an almost perpendicular orientation.

**Figure 3.**
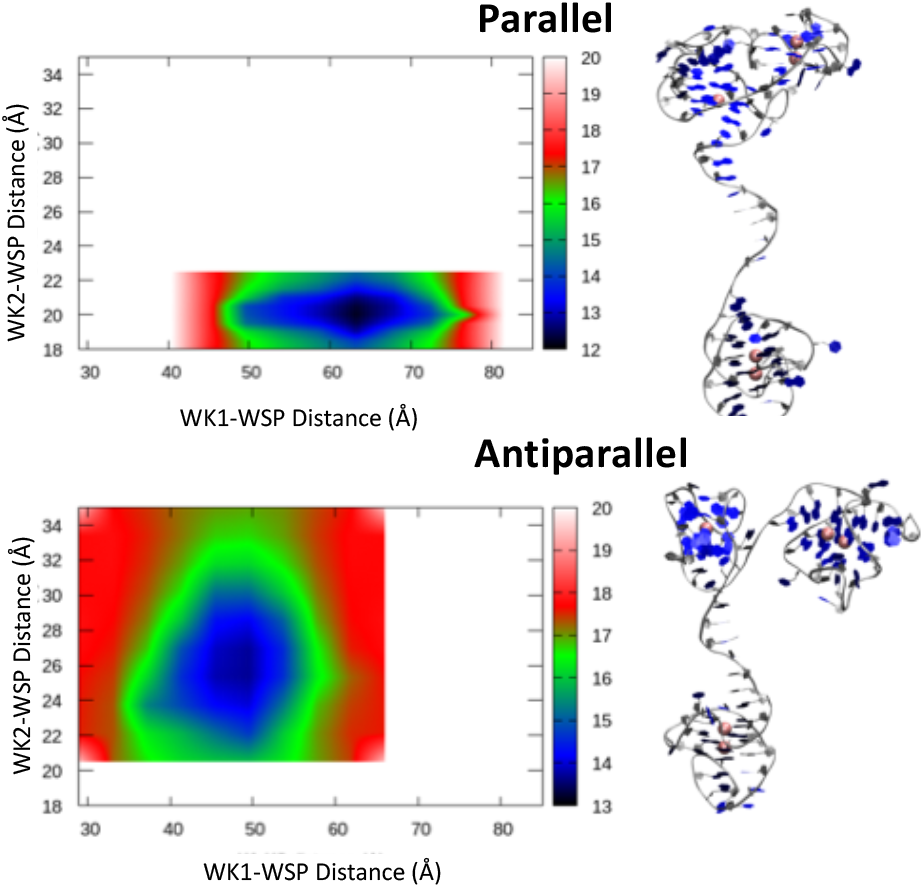
Unbiased free energy profiles (in kcal/mol) along the WK1/WSP and WK2/WSP center of mass distance obtained from the GAMD simulations of the c-*KIT* promoter containing parallel (top) and antiparallel (bottom) WSP. A representative structure corresponding to the free energy minimum is also provided for both conformations.

In addition to the significant differences in the equilibrium distances, we also confirm the higher flexibility of the antiparallel WSP-containing systems. Indeed, the accessible inter-G4 allowed distances are much wider for the antiparallel arrangement, while the parallel WSP results in a sharper free energy landscape. The free energy penalty to disrupt the most favorable arrangement, appears slightly higher for the parallel conformations, of about 8 kcal/mol.

The destabilization of the stacked conformation for the antiparallel WSP is attributable to its less compact structure, which induces non-negligible steric hindrance, impeding the optimal approaching of the two moieties which are instead oriented towards opposite directions. On the contrary, and as already been shown by equilibrium MD, the enhanced flexibility of the antiparallel-containing system leads to a more pronounced folding of the sequence connecting WK1 and WSP, thus slightly approaching the two moieties.

## CONCLUSIONS

G4s are widely occurring structures in DNA, and more generally in nucleic acids, playing essential biological roles, especially related to genome protection and the regulation of gene expression. Despite their rigid core, they can assume a variety of arrangements. Here we have shown that the analysis of the secondary structures of isolated G4s is insufficient to explain their overall preferred structural arrangements. The presence of adjacent G4 units in the sequence of gene promoters leads in fact to the emergence of unexpected tertiary structure arrangements, different from those of the isolated G4 motifs, thus modifying also the properties and roles of the individual components. Such a level of comprehension is up to date rarely achieved for G4s, for which the majority of studies consider only isolated short sequences in solution. Previous studies on the *KIT* promoter based on biophysical experiments provided conflicting results concerning WSP arrangement.^32,37^ This work shed light on the intramolecular interactions of the multiple G quadruplexes in the c-*KIT* promoter, through characterization at atomic scale of the tertiary aggregates, an approach never attempted so far. Using all atom MD simulation and enhanced sampling, for the first time we show that the three G4s in c-*KIT* promoter do not behave independently. A stable and persistent π-stacked aggregate between the terminal (WK2) and the central (WSP) quadruplexes is formed when the latter is folded in a parallel arrangement. On the contrary, when WSP is in its antiparallel conformation the coupling between the three units is lost and the three motifs behave independently. The enhanced stability of the π-stacked aggregate explains the fact that, differently from the case of the isolated units, the ECD spectrum of the whole sequence only shows signatures indicative of a parallel folding.^32^

The atomistic structure of the three adjacent G4s in a gene promoter sequence highlights their cross-talks and interplay, and provides an interpretative framework to the spectroscopic results obtained so-far. Our work also underlies the fact that the behavior of multiple-G4 containing sequences cannot be reduced to the sum of the properties of their individual components, i.e. to the study of their secondary structure only. On the contrary, complex tertiary arrangements take place, significantly modifying the structural and dynamic properties of the DNA sequence. This, in turn, has a crucial role in determining the interaction of specific sequences with regulatory proteins, hence can largely determine the biological role played by a specific G4. Indeed, the sequences corresponding to WK2 and WSP in the c-*KIT* promoter are known to be binding sites for transcription factors such as Sp1 and AP2.^35,36^ Our work contributes to shade a new light on an usually overlooked problem, and bring further exciting perspective for a deeper understanding of the molecular bases of gene regulation. Identifying crucial and complex tertiary structures in G4s will also allow to significantly improve the drug-design strategies aimed at producing G4-stabilizing agents. In particular, the structural complexity of the tertiary arrangements must be used to identify conserved and specific motifs which will enhance the selectivity of the designed agents compared to B-DNA or other G4s.

## METHODOLOGY

All the MD simulations were performed with the NAMD 3 software.^39,40^ The structure of the three G4 were taken from the Protein Data Bank for WK2 (PDB code: 2KYP),^41^ WK1 (PDB code: 2O3M),^33^ and the anti-parallel WSP (PDB code: 6GH0).^36^ The parallel conformation of WSP was generated by homology modeling as detailed in Supplementary Information. The 5’-tail and linker sequences were generated *in silico* and attached to WK2 and WK1, respectively. The full sequence was the one used by Ducani et al.^32^ with a total of 78 residues. The three starting systems (5’-tail+WK2, WSP, linker+WK1) were individually minimized and equilibrated along a 22 ns simulation in water at 300K in NTP, using the ff14SB force field^42^ and bsc1 corrections^43–44^ for DNA backbone dihedrals. Finally, the relaxed structures were connected to each other to generate the full length system after removal of the water and counter-ions used for the equilibration.

The two systems, with WSP parallel or anti-parallel, were then soaked again into a 25Å buffer TIP3P^45^ water box with potassium counter-ions to ensure neutrality. This resulted in systems of ~145*120*150 Å^3^ box size, for a total of up to 240,000 atoms. After minimization and equilibration with decreasing constraints on the DNA backbone, a 900 ns production run in NTP was performed for each system. Simulations were performed using a Langevin thermostat with a 1 ps^-1^ damping frequency to keep the temperature at 300K. Hydrogen Mass Repartioning^46^ was applied to allow the use of a 4 fs time-step. Electrostatics were treated using the Particle Mesh Ewald^47^ with a 9 Å cutoff. The clustering analysis was performed on each MD ensemble based on the RMSD of the 78 DNA residues and using the hierarchical-agglomerative and average-linkage approaches.

In order to enhance the conformational sampling of the full length structures, Gaussian accelerated MD simulations^48^ of 600 ns were performed on both systems with a dihedral potential boost. The threshold energy for adding boost potential was set to its lower bound, and the default value of the limit of the standard deviation σ_D_ was used (6.0 kcal/mol). The 2D free energy maps were generated by reweighting of simulations using the PyReweighting script available on the Miao Lab website (http://miaolab.org/PyReweighting/),^49,50^ taking the distances between the centers of mass of WK2 and WSP, and WKP and WK1 as inputs.

All plots and figures were rendered using the VMD^51^ and gnuplot softwares.

## Supporting information

PDB Structures

PDB structures

## ASSOCIATED CONTENT

### Supporting Information

The Supporting Information is available free of charge on the ACS Publications website.

Time evolution of the root-mean-square-deviation (RMSD), evolution of the population obtained from the clustering, details on the homology model used to reconstruct the parallel arrangement of WSP (ESI.pdf PDF)

Three main cluster poses of the c-*KIT* promoter bearing WSP in parallel arrangement (par_WSP.pdb PDB)

Three main cluster poses of the c-*KIT* promoter bearing WSP in antiparallel arrangement (anti_WSP.pdb PDB)

## AUTHOR INFORMATION

### Author Contributions

The manuscript was written through contributions of all authors.

## ACKNOWLEDGMENT

The authors thank GENCI and Explor computing centers and the Platform P3MB for computational resources. A.M. thanks ANR and CGI for their financial support of this work through Labex SEAM ANR 11 LABX 086, ANR 11 IDEX 05 02. The support of the IdEx “Université Paris 2019” ANR-18-IDEX-0001. A.T. thanks EUROSTART

## TOC Graphic

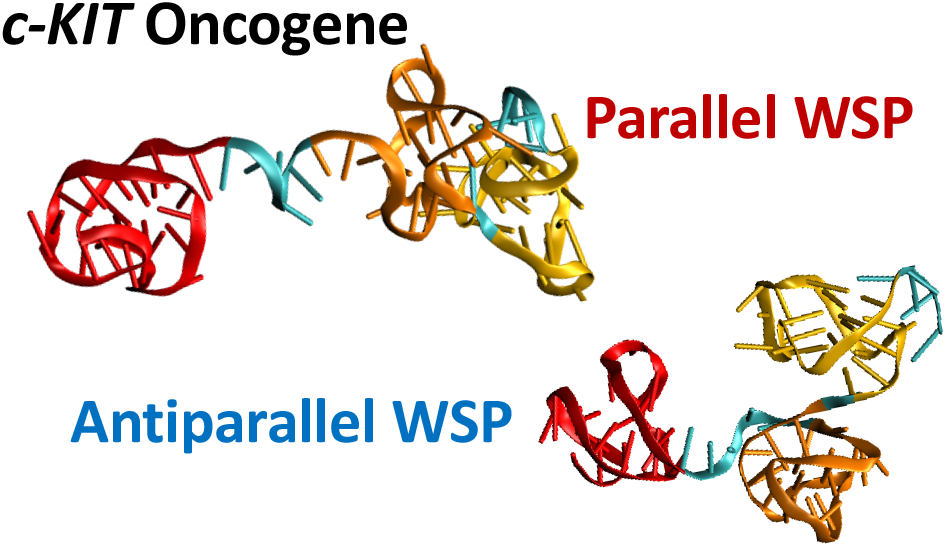

