## Supplementary Information for "Predicting the Three-dimensional Structure of the c-*KIT* Oncogene Promoter and the Dynamics of Strongly Coupled Guanine-Quadruplexes"

Electronic Supplementary Information for:

a) Université de Lorraine and CNRS, UMR 7019 LPCT, F-54000 Nancy, France. b) Università di Palermo, Department of Biological, Chemical and Pharmaceutical Sciences and Technologies Viale delle Scienze, Edificio 17, 90128 Palermo, Italy. c) Department of Medical Biochemistry and Biophysics, Karolinska Institutet, Stockholm 171 65, Sweden, d) Moligo Technologies AB, Banvaktsvägen 20, 171 48 Solna, Sweden e) Université de Lorraine and CNRS, UMR 7039 CRAN, F-54000 Nancy, France. f) Université Paris Cité and CNRS, ITODYS, F-75006 Paris, France.

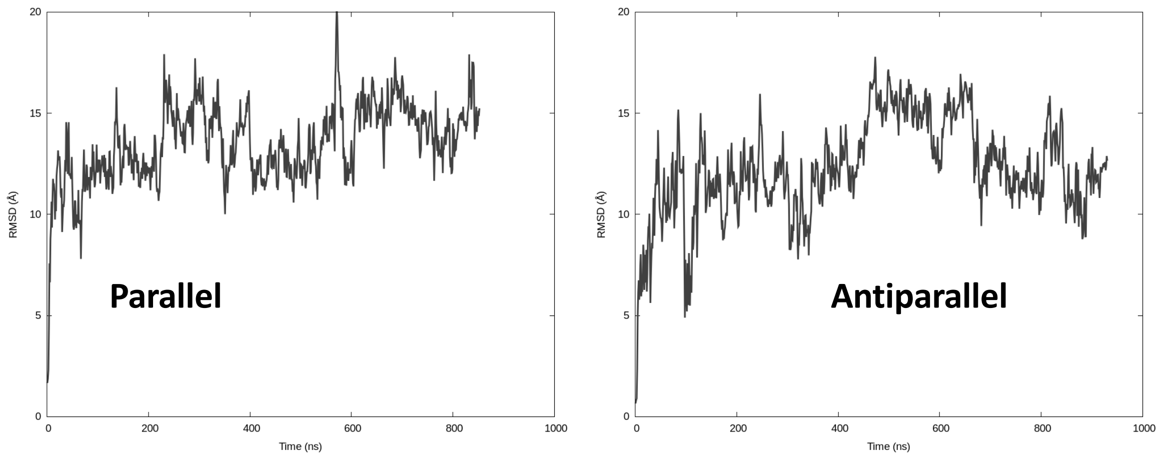


Figure S1) Time evolution of the RMSD of the whole *c-KIT* promoter with WSP in parallel (left) and antiparallel (right) arrangement, respectively.

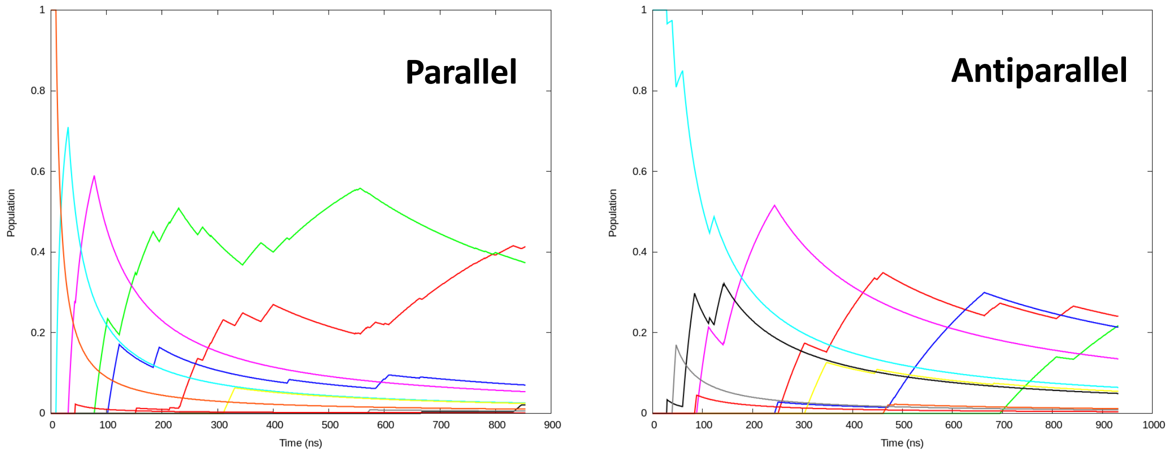


Figure S2) Evolution of the population of the ten most important cluster for the *c-KIT* promoter featuring WSP in parallel (right) and antiparallel (left) conformation, respectively.

### Details on the homology modeling to retrieve initial parallel WSP arrangement

*1) Structure search in the PDB database*

The homology search pursued here follows the same methodology used in a previous work.[1] The target WSP sequence is composed of 21 nucleotides, 17 of which form the G-quadruplex core. This size necessitates a search by motif in the PDB database, with the parameter PROSITE. However, by using only this parameter we obtain only one hit, i.e. PDB:139D, however this entry corresponds to a parallelel G4 composed by four distinct strands. Thus, the absence of connecting loops precludes the use of this structure. Then two other searches were performed, the first using the keyword "quadruplex", and the second one using structure similarity based on the PDB:1KF1 structure together with the Relaxed parameter. The top five structures issued from each search were used to perform a refinement.

*2) Refinement by sequence similarity*

M-Coffee server[2] is used to perform the multiple sequence alignment between WSP and the ten top sequences obtained by the structure search. The sequences only share limited similarities have little similarity (Figure S3). Consequently, the alignment result does not always match the guanines forming quartets. While PDB:6T51 appears to be the closest to WSP, the further verification of this structure has highlighted the presence of bulge in the tetrad, therefore, this sequence is discarded. Consistently, PDB:24NY is also discarded for similar reasons.

```
WSP      G--GC-GAGGAGGGGCGTGGCCG----------G
1XAV     T--GAGGGTGGGTAGGGTGGGTA----------A
6H1K     G---G-GAGGCGTGGCCTGGGCGGGACTGGG--G
3CDM     T--AG-GGTTAG-GGTTAGGGTTAG----G---G
5DWX     G-----GGTGGGTGGGT-GGGTTAG----CGTTA
5DWW     G-----GGTGGGTGGGTTGGGTTAG----CGTTA
6ISW     ---AG-GGTTAG-GGTXAGGGTTAG----G---G
2LD8     T--AG-GGTTAG-GGTTAGGGTTAG----G---G
2N4Y     ----C-TGGGCGGGACTGGGGAGTG-----G--T
6T51     AGGGC-GGTGTGGGAATAGGGA-----------A
cons                *      **
```

Figure S3) Multiple sequence alignment between WSP and the ten top hits issued from structure search in the pdb database. Note the rather low similarity between the sequences.

*3) Refinement by structure comparison*

After manually positioning the structural elements of each G4 we observe the presence of nucleotides not involved in the formation of the G4 at the 5' and 3' positions.  Besides, the structural comparison shows a large difference in the size and extension of the connecting loops (Figure S4). The only structures presenting loops of three nucleotides, thus analog to WSP, are PDB:3CDM, PDB:6ISW, and PDB:2LD8. Furthermore, the loops in these structures share the same sequence: TTA, except for PDB:6ISW (Figure S4).

```
WSP     ---GG-CGA--------GG-AGGGG-CGTGG-CCGG-----
1XAV    TGAGGGT----------GGGTA-GGGT--GGGTAA------
6H1K    ---GGGAGGCGTGGCCTGGGC--GGGACTGGGG--------
3CDM    -TAGGGTTA--------GGGTTAGGGTTAGGG---------
5DWX    ---GGGT----------GGGT--GGGT--GGGTTAGCGTTA
5DWW    ---GGGT----------GGGT--GGGTT-GGGTTAGCGTTA
6ISW    --AGGGTTA--------GGGTXAGGGTTAGGG---------
2LD8    -TAGGGTTA--------GGGTTAGGGTTAGGG---------
```
Figure S4) Structural comparison between WSP and the top hits G4 from the PDB database.

*4) Construction of the parallel WSP G-quadruplex structure*

The final structure was obtained considering PDB:3CDM and PDB:2LD8. Notably Maestro software[3] was used to remove the third tetrad (which is absent in WSP) and eliminate the 5' T and add the 3' CCGG cap. Instead, A in 5' to the first tetrad is maintained because it corresponds to the nucleotide linking WK2 and WSP in the *c-KIT* sequence. An intermediate structure has been generated with UCSF Chimera[4] as a pdb file using a nomenclature compatible with the Amber force field. Finally, the nucleotides present in the loops were modified with the Mutagenesis tool of PyMol.[5]


(1) Miclot, T.; Hognon, C.; Bignon, E.; Terenzi, A.; Marazzi, M.; Barone, G.; Monari, A. Structure and Dynamics of RNA Guanine Quadruplexes in SARS-CoV-2 Genome. Original Strategies against Emerging Viruses. *J. Phys. Chem. Lett.* **2021**, *12* (42), 10277–10283. https://doi.org/10.1021/acs.jpclett.1c03071.

(2) Moretti, S.; Armougom, F.; Wallace, I. M.; Higgins, D. G.; Jongeneel, C. V.; Notredame, C. The M-Coffee Web Server: A Meta-Method for Computing Multiple Sequence Alignments by Combining Alternative Alignment Methods. *Nucleic Acids Res.* **2007**, *35* (SUPPL.2). https://doi.org/10.1093/nar/gkm333.

(3) Schrödinger, LLC, New York, N. Schrödinger Release 2021-1: Maestro. 2021.

(4) Pettersen, E. F.; Goddard, T. D.; Huang, C. C.; Couch, G. S.; Greenblatt, D. M.; Meng, E. C.; Ferrin, T. E. UCSF Chimera - A Visualization System for Exploratory Research and Analysis. *J. Comput. Chem.* **2004**, *25* (13), 1605–1612. https://doi.org/10.1002/jcc.20084.

(5) Schrödinger, LLC. *The {PyMOL} Molecular Graphics System, Version~1.8*; 2015.
